# Anti-androgens induce Rab11a-exosome secretion in prostate cancer by suppressing amino acid-sensitive PAT4-mTORC1 signalling

**DOI:** 10.1101/2020.09.18.300137

**Authors:** Kristie McCormick, Poohrawind Sanitt, Shih-Jung Fan, John D Mason, Adrian L Harris, Freddie C Hamdy, Clare Verrill, Richard J Bryant, Deborah C I Goberdhan

## Abstract

Advanced prostate cancer is typically treated with anti-androgens to reduce cancer growth, but patients almost inevitably develop treatment resistance and castration-resistant disease. Recently, extracellular vesicles known as exosomes, which are secreted from the endosomal compartments in which they are formed, have been implicated in drug resistance mechanisms. Here we investigate whether growth regulation by the amino acid-dependent kinase complex, mechanistic Target of Rapamycin Complex 1 (mTORC1), and associated extracellular vesicle secretion might be involved in the adaptive responses to anti-androgens. We show that expression and intracellular localisation of the glutamine-sensing PAT4 (SLC36A4) amino acid transporter is increased in malignant versus benign prostatic tissue, mirroring earlier *in vivo* fly studies suggesting that these transporters are more effective at promoting growth from internal versus cell surface membranes. Furthermore, androgens induce PAT4 expression in prostate cancer cell lines and PAT4 is required for a proportion of androgen-stimulated mTORC1 activation and growth. Consistent with previous studies in other cancer cell lines, we find that glutamine depletion, PAT4 knockdown and mTORC1 inhibition all independently increase the production of a specific exosome subtype, Rab11a-exosomes, which has recently been implicated in pro-tumorigenic signalling responses to mTORC1 inhibition. Furthermore, we show that these exosomes are also induced by anti-androgens. We hypothesise that the uptake of Rab11a-exosomes by cells with higher PAT4 levels could provide a growth-promoting boost, enabling them to out-compete others with lower PAT4 expression, resulting in tumours that are more resistant to nutrient-deprivation and anti-androgen treatment.

## Introduction

Prostate cancer (PCa) is the most common solid organ malignancy in men and the second most common cause of male cancer-related death (Cancer Research UK, 2016). Androgen deprivation therapy (ADT) is a major modality used to treat locally advanced and metastatic PCa, but a ‘castration-resistant’ form of PCa (CRPCa) typically develops, which is usually fatal within approximately two years (Tannock et al, 2004). Identifying the mechanisms by which this CRPCa state emerges is an area of intense interest (Nakazawa et al, 2017). In beniognl and malignant prostate cells, androgens have been shown to promote mechanistic Target of Rapamycin (mTORC1) activity (Xu et al., 2006; Rai et al., 2010; Wu et al., 2010; Tsouko et al., 2014), an integrator of microenvironmental growth factor and nutritional signals (Goberdhan et al., 2016). A key way in which androgen signalling modulates mTORC1 activity is by increasing the expression of proteins that enhance nutrient availability and cellular metabolism (Waghray et al., 2001; Nelson et al., 2002; Deprimo et al., 2002; Velasco et al., 2004; Sharifi et al., 2012). For instance, several transporters belonging to the SLC family of proteins are upregulated by androgens (Xu et al., 2006) and have been shown to mediate the effects of androgens on PCa cell growth by enhancing mTORC1 activity (White et al., 2017).

Our previous studies have identified the SLC family member Proton-assisted Amino-acid Transporter 4 (PAT4/SLC36A4) as a glutamine-sensitive amino acid transporter and positive regulator of mTORC1 signalling and cell proliferation (Heublein et al., 2010; Fan et al., 2016). Furthermore, we demonstrated that high levels of PAT4 are associated with diminished relapse-free survival after colorectal cancer surgery. Consistent with this, PAT4 promotes HCT116 colorectal cancer growth in both *in vitro* cell culture and in *in vivo* xenograft tumour models (Fan et al., 2016).

Although the acquisition of genetic mutations can drive therapeutic drug resistance mechanisms, another route by which resistance can emerge is through cell-cell communication within the heterogeneous tumour and with its stroma (Cheng et al, 2020). One evolutionarily conserved mechanism of intercellular signalling involves the transfer of extracellular vesicles (EVs) between cells (Meldolesi, 2018). EVs can transfer ligands, receptors, cytoplasmic proteins, nucleic acids and metabolites between cells as part of normal homeostasis. However, EVs have also been implicated in the oncogenic reprogramming of cells within the tumour microenvironment and the pathogenesis of numerous malignancies (Hoshino et al., 2015; Melo et al., 2015), including PCa (Zhao et al., 2016). For example, EVs secreted from PCa cells have been shown to promote cell proliferation (Soekmadji et al., 2016), angiogenesis (Naito et al., 2017), metastasis (Bijnsdorp et al., 2013), immune evasion (Lundholm et al., 2014), and drug resistance (Corcoran et al., 2012).

The cargos of secreted EVs are critical to their function. EV composition must therefore be under tight regulation. For example, various stress stimuli, such as hypoxia, acidosis, starvation and anti-cancer therapies have been shown to modulate EV content (Parolini et al., 2009; Kucharzewska et al., 2013; Azmi et al., 2013). We recently demonstrated that the mTORC1 signalling axis suppresses the secretion of a novel population of EVs, termed Rab11a-exosomes (Fan et al., 2020; reviewed by van Niel and Théry, 2020). Exosomes are a type of EV made by the inward budding of the limiting membrane of endocytic compartments, which have classically been thought to be late endosomes. However, Rab11a-exosomes, which are found in human and *Drosophila* cells, are made in compartments marked by the recycling endosomal marker Rab11a, and some of these vesicles are Rab11a-positive. They are preferentially secreted from tumour cells following either glutamine depletion or pharmacological inhibition of mTORC1 activity and they exhibit enhanced pro-tumorigenic functions in target cells, which may mediate adaptive responses of tumours to microenvironmental stress.

Given the previously described role of androgens in controlling mTORC1 activity, we hypothesized that androgen deprivation might also promote the secretion of Rab11a-marked EVs from PCa cells. Here we show that high levels of PAT4 expression and its increased intracellular localisation are associated with poor outcome in PCa, and that androgens promote mTORC1 signalling in prostate cancer cells partly through the upregulation of PAT4 expression. Furthermore, anti-androgen treatment induces Rab11a-exosome secretion, suggesting a mechanism by which tumour cells might respond and adapt to such treatments through exosome signalling, thereby promoting the growth of PCa cells with high PAT4 levels and enhanced treatment resistance.

## Results

### PAT4 protein is increased and its distribution shifts to intracellular compartments in malignant versus benign prostatic tissue

To test whether PAT4 expression is altered in PCa, we performed immunohistochemistry with an anti-PAT4 antibody using a tissue microarray (TMA) containing tissue samples from 137 PCa patients and 44 individuals with benign prostatic hyperplasia (BPH) (Figure 1a). Immunohistochemistry revealed variable PAT4 subcellular localisation in the plasma membrane, cytoplasm, and nuclei in both the BPH and PCa samples. In BPH, the mean cytoplasmic immunoreactive signal measured by the standardised immunoreactive score (IRS) system was 2.41 ±0.26 (mean ± SEM), compared to 6.54 ± 0.33 in PCa, whereas the mean membrane IRS in BPH was 4.66 ± 0.47 versus 3.26 ± 0.36 in PCa, indicating that there is significantly more PAT4 at the plasma membrane in BPH when compared to PCa (p<0.02) and significantly more PAT4 in the cytoplasm of PCa when compared to BPH (p<0.001). This is consistent with previous studies, suggesting that oncogenic PI3K signalling increases the intracellular localisation and growth-promoting properties of PAT amino acid transporters (Ögmundsdóttir et al., 2012). Overall, PAT4 expression was higher in the malignant prostatic epithelium, when compared to benign glands (Figure 1b).

**Figure 1.**
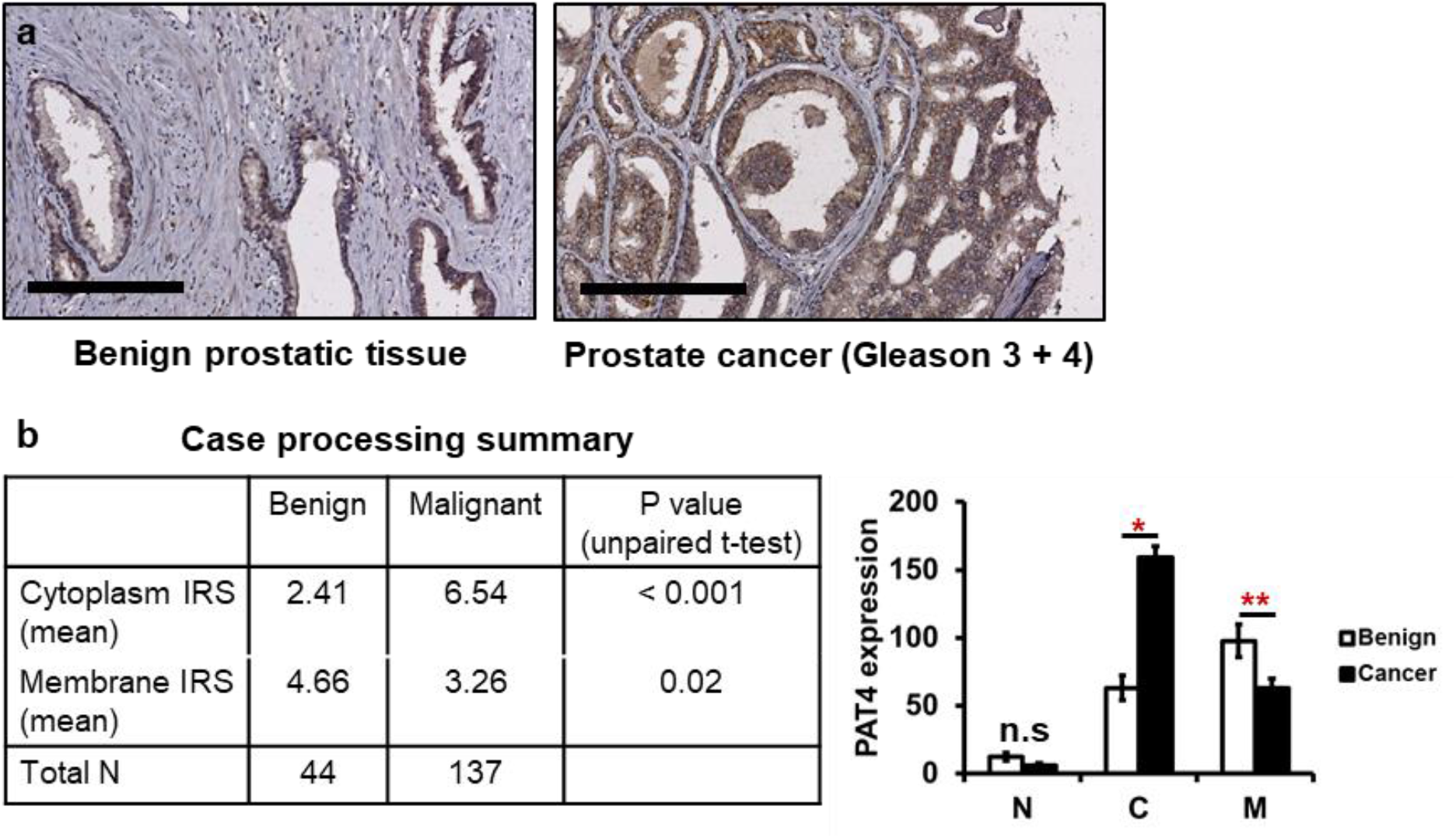
PAT4 expression and localisation to intracellular compartments is increased in prostate cancer. (**a**) Tissue samples in a microarray from 137 patients with primary prostate carcinoma (Gleason grades three and four) and 44 individuals with benign prostatic hyperplasia were stained with an anti-PAT4 antibody. (**b**) Bar charts show the nuclear (N), cytoplasmic (C) and membranous (M) expression levels of PAT4 (arbitrary units) in PCa and BPH tissue. The scale bars in (**a**) are 200 μm. ns > 0.05; \**P* < 0.05; \*\**P* < 0.01.

### Androgen-induced mTORC1 activation and cell growth are partially mediated by PAT4 signalling

Since androgens promote expression of several SLC amino acid transporters, we examined whether androgens also regulate PAT4 expression. The effects of androgens on *PAT4* mRNA were assessed in androgen-sensitive VCaP and LNCaP prostate cancer cell lines. Treatment of VCaP cells with the androgen dihydrotestosterone (DHT) for 48 hours significantly increased the expression of *PAT4* mRNA, mirroring the increase observed in the prostate cancer biomarker, prostate-specific antigen (PSA; Figure 2a). Expression of the glutamine-sensitive transporter, which is known to be upregulated in PCa (White et al., 2017), was also increased following DHT treatment, but expression of the other ubiquitously expressed PAT transporter, PAT1, was unaffected. In addition, DHT significantly increased PAT4 protein levels in VCaP cells after 48 h (Figure 2b), consistent with the results observed for *PAT4* mRNA. In contrast, in LNCaP cells, PAT4 mRNA levels were unaffected by DHT treatment, despite androgen stimulation resulting in increased PSA and SLC1A5 expression (Figure 2a).

**Figure 2.**
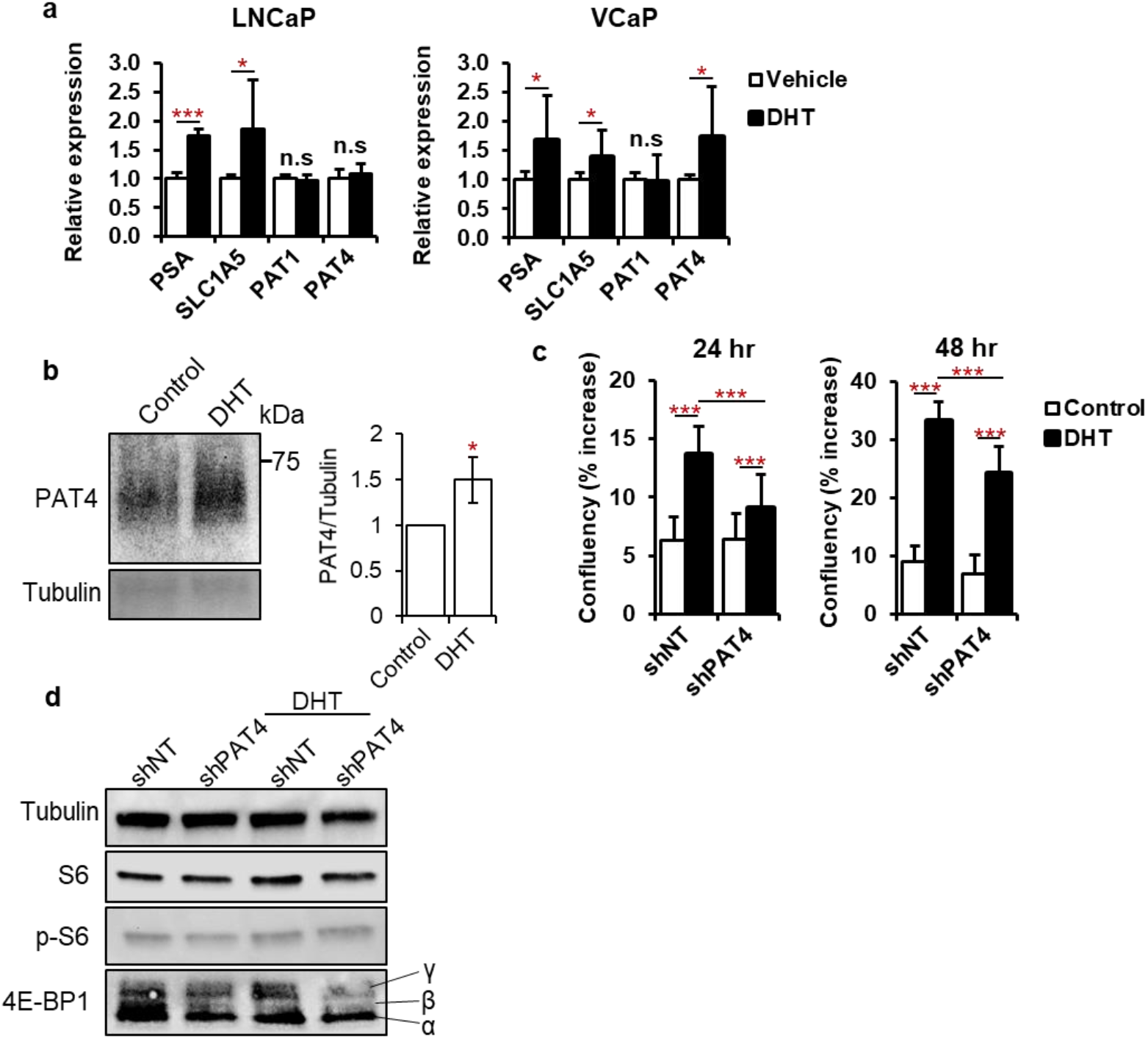
Androgen-induced mTORC1 activation and cell growth is partially mediated by PAT4. (**a**) The mRNA levels of the PAT transporters, and androgen-sensitive SLC1A5 transporter and prostate-specific antigen (PSA) in VCaP and LNCaP prostate cancer cell lines, as measured by qRT-PCR, in the presence or absence of dihydrotestosterone (DHT). (**b**) Western blot analysis of cell lysates from VCaP cells with or without DHT treatment, using an anti-PAT4 and anti-tubulin antibody. (**c**) Growth of VCaP cells (measured as percentage confluency) in the presence or absence of DHT, and treated either with a *P∠47W*-shRNA (shPAT4) or a non-targeting shRNA control (shNT). (**d**) Western blot analysis of cell lysates for mTORC1 downstream targets, phospho-S6 (p-S6) and the γ-phosphorylated form of 4E-BP1 (top band, which increases in response to DHT in shNT-treated cells). ns > 0.05; \**P* < 0.05; \*\**P* < 0.01; \*\*\**P* < 0.001

To determine the role of PAT4 in androgen-induced growth, VCaP cells were transduced with constructs either expressing a short-hairpin RNA targeting PAT4 (shPAT4), or a non-targeting control (shNT), and cultured in the presence or absence of DHT for 48 h, and the effects on cell growth were assessed. In control shNT-treated cells, addition of DHT significantly increased cell growth at both 24 and 48 h (Figure 2c), consistent with the reported effects of androgens on these cells (Pretorius et al, 2016). Cells also increased growth after DHT stimulation following *PAT4* knockdown, but the response was significantly decreased compared to controls, suggesting that PAT4 expression is involved in androgen-induced growth stimulation. Treatment with DHT stimulated mTORC1 activity (Figure 2d), as evidenced by the increase in phospho-S6 (pS6) and hyperphosphorylated 4E-BP1, consistent with findings reported in the published literature (Xu et al, 2006). This effect was not observed in PAT4-deficient cells, indicating that, as with cell growth, PAT4 is necessary to permit mTORC1 activation in response to androgens. In addition, *PAT4* knockdown also inhibited basal mTORC1 activity, suggesting that PAT4 regulates mTORC1 signalling in the presence and absence of androgens.

### Glutamine depletion and mTORC1 inhibition induce the secretion of Rab11a-exosomes in prostate cancer cell lines

We have previously shown that glutamine depletion increases the secretion of a Rab11a-marked population of exosomes from LNCaP cells (Fan et al., 2020). To test the role of PAT4 in the production of Rab11a-exosomes in other PCa cells, we first characterised the effects of glutamine depletion on VCaP and C4-2B cells (a bone metastatic subline of LNCaP cells), using an equivalent culture method and exosome isolation protocol. As expected, following glutamine depletion, the phosphorylation status of two well-characterised downstream mTORC1 targets, 4E-Binding Protein 1 (4E-BP1) and S6, was reduced in the VCaP cell line (Figure 3a). Interestingly, however, no difference in phosphorylation of either target was detected in C4-2B cells after 24 h (Supplementary Figure 2). Nanoparticle Tracking Analysis (NTA) of EV preparations revealed that EV numbers were similar to control values after glutamine depletion, and that EV size distribution was unaltered.

**Figure 3.**
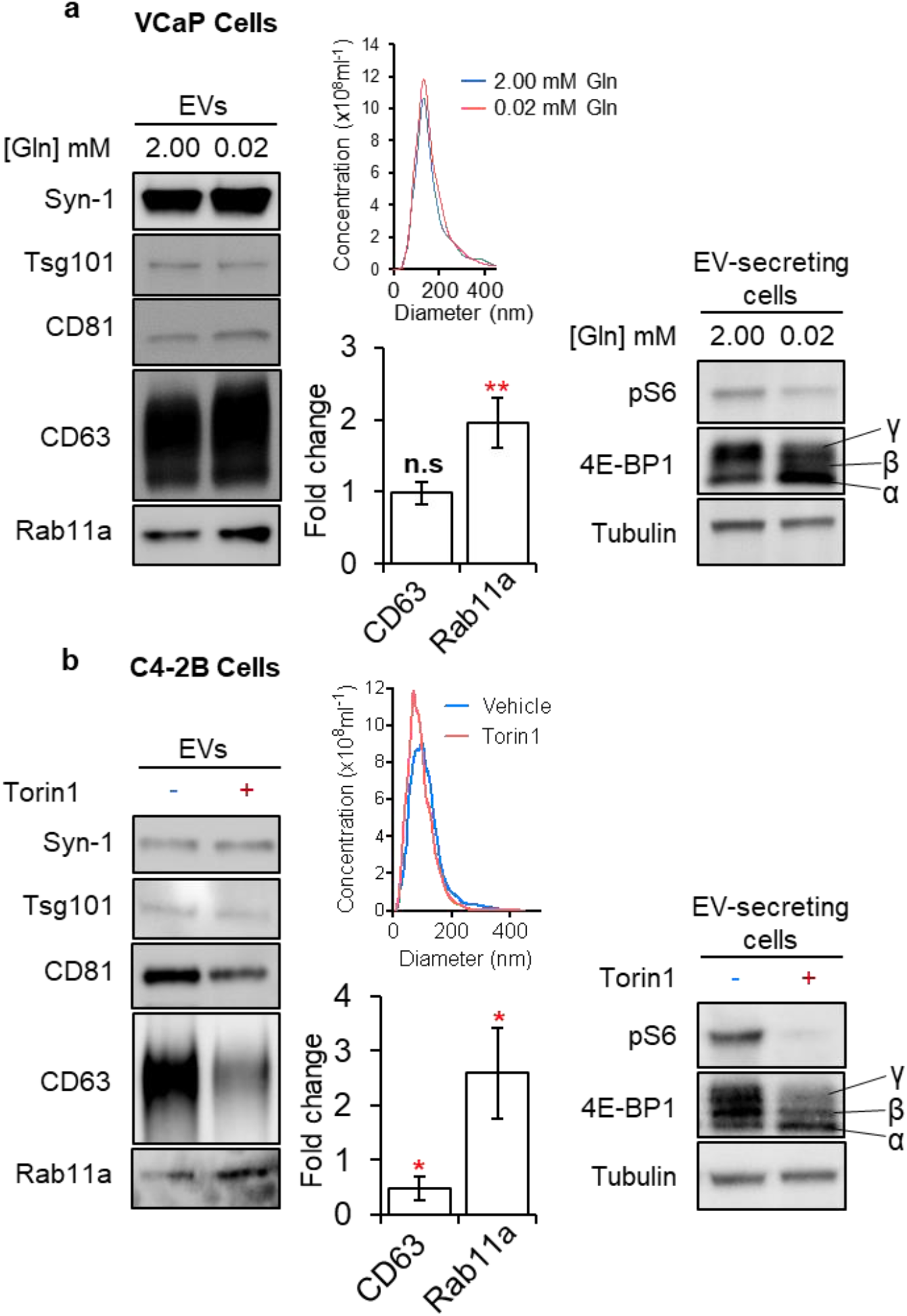
Glutamine depletion and mTORC1 inhibition induce the secretion of Rab11a-exosomes in prostate cancer cell lines. (**a**) Western blot analysis of cell lysates and extracellular vesicles (EVs) from VCaP cells cultured in glutamine-replete (2.00 mM) and glutamine-depleted (0.02 mM) medium for 24 h. Levels of phospho-S6 and the γ-phosphorylated form of 4E-BP1 in cell lysates were determined to assess mTORC1 activity. EVs were loaded according to cell lysate protein levels to compare secretion of putative exosome markers on a per cell basis. Bar charts are from three biological repeats. (**b**) Western blot analysis of cell lysates and extracellular vesicles from C4-2B cells with or without Torin1 treatment for 24 h. The activity of mTORC1 was assessed via phosphorylation of the S6 (pS6) and 4E-BP1, as described above. Bar charts present data from three biological repeats. The number of EVs in each condition was determined using NanoSight analysis. ns > 0.05; \**P* < 0.05; \*\**P* < 0.01

Glutamine depletion of VCaP cells significantly increased the levels of secretion of Rab11a (Figure 3a), while other exosome markers were unchanged, findings mirroring those reported for LNCaP cells (Fan et al., 2020). Markers unaffected by glutamine depletion included CD81, CD63, Syntenin-1, Alix, and Tsg101. In contrast, no change in EV content was observed for C4-2B cells following glutamine depletion (Supplementary Figure 2). However, treatment of C4-2B cells with the mTOR-specific inhibitor, Torin1, did strongly increase the secretion of Rab11a in EVs, while CD63 content significantly decreased (Figure 3b), as has been observed in several other cancer cell lines (Fan et al., 2020). This suggests that these cells are insensitive to glutamine, but still regulate exosome secretion through mTORC1.

### *PAT4* knockdown induces a switch to secretion of Rab11-exosomes in VCaP cells

Since glutamine-sensing PAT4 regulates mTORC1 activity in VCaP cells, we tested whether *PAT4* knockdown also affects EV release from these cells. EVs were collected for 24 h on the sixth day of knockdown. Cellular levels of hyperphosphorylated 4E-BP1 and pS6 were reduced following *PAT4* knockdown (Figure 4), confirming our earlier findings. The knockdown also induced a strong increase in Rab11a and a slight decrease in CD63 within EV preparations. These findings are consistent with the induction of an mTORC1-regulated switch in the balance of exosome secretion from late endosomal to recycling endosomal compartments.

**Figure 4.**
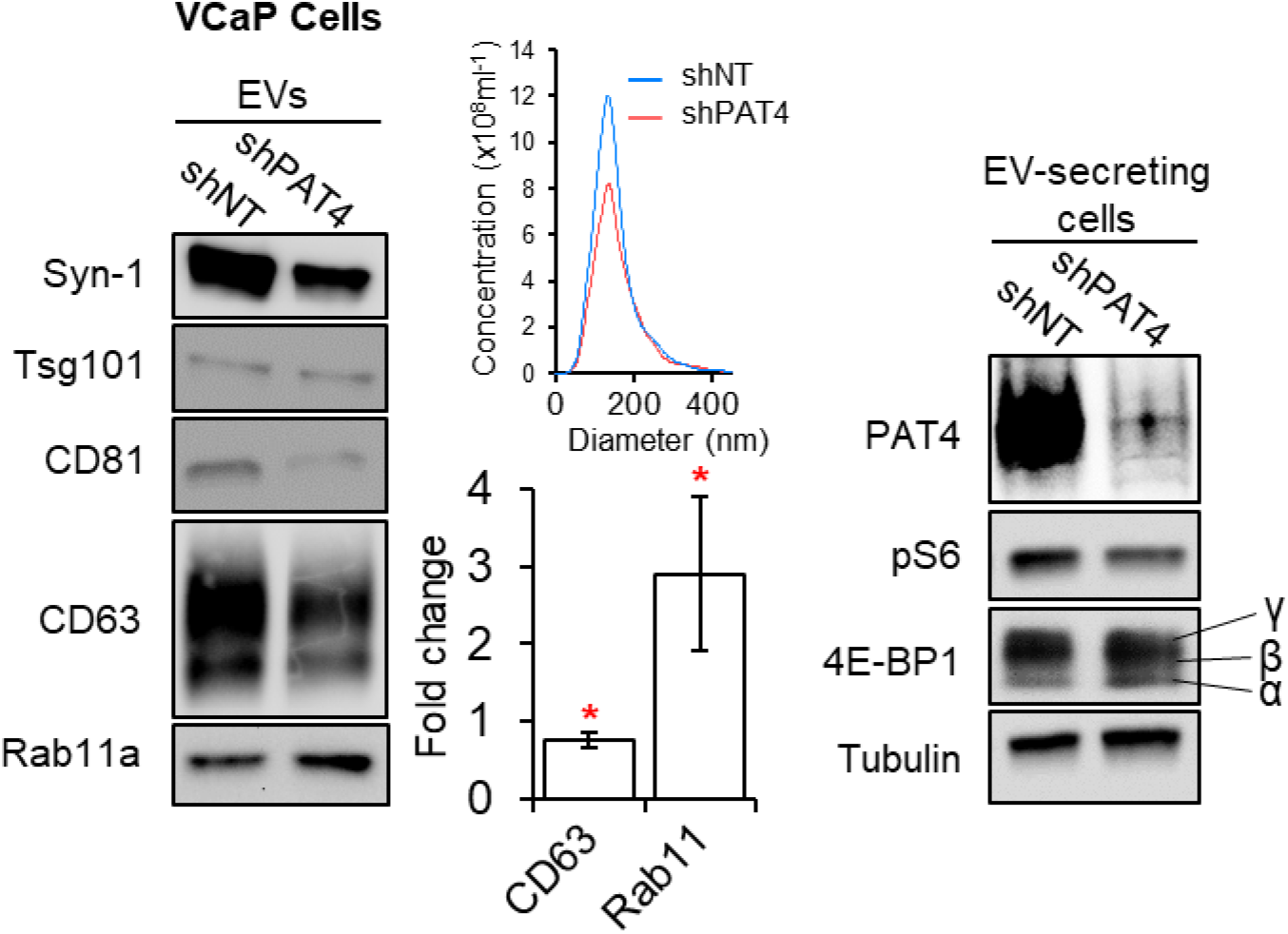
*PAT4* knockdown induces a switch to secretion of Rab11a-exosomes in VCaP cells. Western blot analysis of cell lysates and EVs from VCaP cells following transfection with *PAT4* (shPAT4) or non-targeting (shNT) shRNA constructs for five days. The activity of mTORC1 was assessed via phosphorylation of S6 and 4E-BP1, in addition to levels of PAT4. EV lysates were probed for putative exosome proteins. Bar charts present data from three biological repeats. \**P* < 0.05

### Androgen-deprivation induces a switch to secretion of Rab11a-exosomes in VCaP cells

As androgen signalling upregulates PAT4 expression and knockdown of *PAT4* induces the release of Rab11a-marked EVs, we hypothesised that androgen-deprivation might drive similar changes in EV cargo. To test this, we cultured VCaP cells in medium supplemented with 1% pre-spun charcoal-stripped serum (PS-CSS) plus DHT and treated the cells with either vehicle (i.e. DMSO) or the anti-androgen bicalutamide for 24 hours. Different EV culture conditions were utilised in this case, as it proved difficult to modify mTORC1 activity in VCaP cells using bicalutamide when the medium was supplemented with ITS. Using these alternative methods, we observed that growth factor signalling was maintained in VCaP cells over the 24-hour EV collection period and we were able to successfully isolate EVs from cells cultured in these conditions. Any remaining EVs present in the pre-spun CSS had minimal influence on the protein signal detected from the EVs isolated from VCaP cells (Supplementary Figure 3).

As expected, following bicalutamide treatment, cellular levels of hyper-phosphorylated 4E-BP1 and pS6 were reduced in the VCaP cell line after 24 h; cellular expression of exosome proteins was unaffected (Figure 5). NTA of EV preparations revealed that EV numbers were similar to control values after androgen deprivation, and that EV size distribution was unaltered. However, bicalutamide treatment selectively increased the secretion of Rab11a by nearly three-fold (Figure 5), closely paralleling the findings observed in these cells with glutamine depletion and *PAT4* knockdown. Therefore, bicalutamide induces a switch in exosome secretion in PCa cells that is similar to the stress-induced switch we have previously seen in colorectal cancer cells, which drives pro-tumorigenic effects and potential adaptation in target cells.

**Figure 5.**
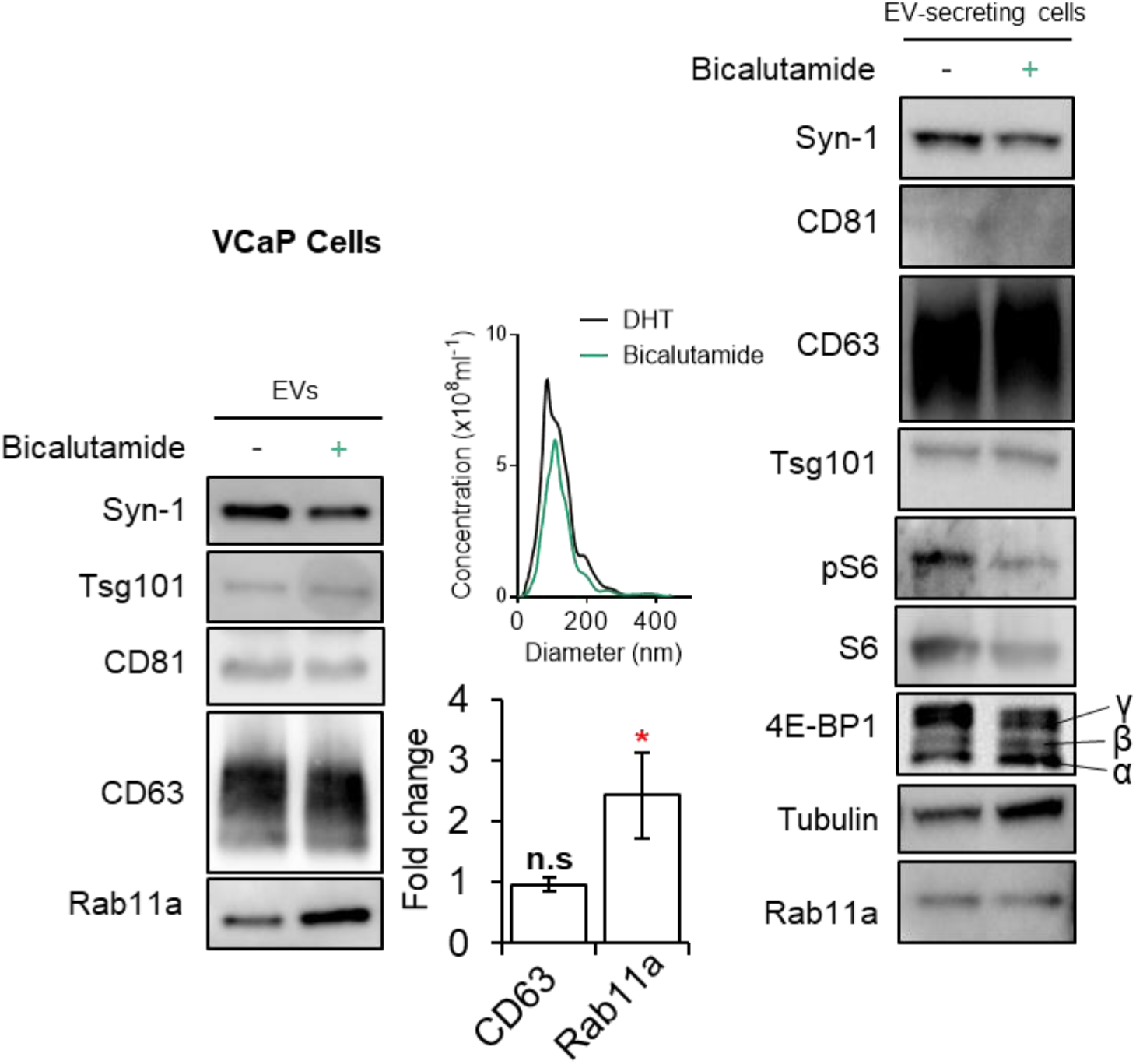
Androgen-deprivation induces a switch to secretion of Rab11a-exosomes from VCaP cells. Western blot analysis of cell lysates and EVs from VCaP cells in the presence of DHT, with and without bicalutamide treatment (Bic), for 24 h. The activity of mTORC1 was assessed in cell lysates via phosphorylation of S6 and 4E-BP1. EV lysates were probed for putative exosome proteins. Bar charts present data from three biological repeats. The number of EVs in each condition was determined using NanoSight analysis. \**P* < 0.05

## Discussion

Advanced PCa that has metastasised outside of the prostate gland to distant sites such as bone or other organs is currently incurable. Although treatment options, such as ADT, next-generation ADT and chemotherapy, can reduce tumour progression for several years, treatment relapse usually occurs, and subsequent treatment options are palliative. Many patients with metastatic PCa develop treatment resistance to ADT, termed CRPCa, within two years. We have used *in vitro* models to examinewhether exosomes might be involved in the develop,ment of CRPCa, focussing in particular on the Rab11a-exosome subtype, which we have recently shown to be produced in response to glutamine depletion and mTORC1 inhibition in several cancer cell lines, resulting in pro-tumorigenic effects (Fan et al., 2020).

We demonstrate that mTORC1 signalling is regulated by the amino acid-sensing transporter PAT4 in VCaP cells. Expression of this transporter is partly controlled by androgens and is elevated in PCa, when compared to BPH tissue. The anti-androgen bicalutamide suppresses mTORC1 signalling, in part due to its effects on PAT4, and this stimulates the production of Rab11a-exosomes. We propose that this mechanism may be involved in selecting cells with high PAT4 levels that are more resistant to treatments, such as ADT.

### PAT4 expression and intracellular localisation are increased in malignant PCa tissue

We have previously shown that high levels of PAT4 expression correlate with reducedrelapse-free survival after colorectal cancer surgery (Fan et al., 2016). Rather than activating mTORC1 by transporting amino acids, *in vivo* analysis in *Drosophila* combined with transport studies has highlighted potential roles for PAT transporters as transceptors; such proteins look like transporters, but actually act like receptors (Goberdhan et al., 2005; Pillai and Meredith, 2011). Furthermore, the similarities between the PAT (SLC36) and sodium-coupled neutral amino acid transporter (SNAT; SLC38) family raise the idea that these transporters may share a similar amino acid-sensing mechanism (reviewed in Fan and Goberdhan, 2018).

Multiple studies have indicated that such amino acid sensors regulate mTORC1 signalling from intracellular compartments. For example, PAT1 (SLC36A1) and SLC38A9 (Ögmundsdóttir et al., 2012; Wang et al., 2015; Rebsamen et al., 2015) control mTORC1 from the surface of late endosomal and lysosomal compartments. More recently, other amino acid transporters, including PAT4, have been shown to regulate mTORC1 signalling from the Golgi (Fan et al., 2016; Krokowski et al., 2017; Tripathi et al., 2019; Morotti et al., 2020; Goberdhan et al., 2016). PAT4 is one of these. It has also been reported to traffic through the recycling endosomal system, but it is unclear how this transporter pool might be involved in mTORC1 control (Matsui et al., 2013). We have previously demonstrated that PATs localise to intracellular compartments in response to oncogenic signalling in a *Drosophila in vivo* model. This effect was coupled with an increase in the growth-promoting properties of the PATs, supporting the model in which amino acid sensors become more effective at promoting growth from inside cells (Ögmundsdóttir et al., 2012). Our analysis of PAT4 expression in PCa now reveals for the first time a correlation between intracellular PAT localisation and malignancy. These findings raise the intriguing possibility that oncogenic signalling in the prostate gland may drive PAT4 intracellularly, as previously observed in *Drosophila*, although the effects of PAT4 relocalisation on prostate cancer cell growth are yet to be explored.

### PAT4 regulates mTORC1 signalling in prostate cells and its expression is suppressed by anti-androgens

We observed that PAT4 activates mTORC1 signalling in androgen-sensitive VCaP cells, consistent with previous findings in HCT116 colorectal cancer cells (Fan et al., 2016). In VCaP cells, PAT4 expression is regulated by androgens, mirroring the effects on the glutamine transporter SLC1A5 (White et al, 2017). Importantly, androgen-stimulated growth is partly mediated by PAT4, and treatment with anti-androgens, glutamine depletion or knockdown of *PAT4* suppress growth and mTORC1 signalling.

We have previously demonstrated that glutamine depletion reduces mTORC1 signalling in LNCaP cells. However, here we found that C4-2B cells, which are metastatic derivatives of LNCaP cells, are glutamine-insensitive and therefore may regulate mTORC1 via alternative mechanisms, independently of PAT4. Nonetheless, our clinical data indicate that increased PAT4 expression is common in PCa, suggesting that it may frequently be involved in promoting mTORC1-regulated growth in this malignancy.

### Rab11a-exosome secretion is induced in prostate cancer cell lines following anti-androgen treatment and mTORC1 inhibition

Inhibition of mTORC1 signalling in LNCaP cells induces a switch in exosome secretion with increased release of Rab11a-exosomes (Fan et al., 2020). We have shown that this is also the case for VCaP and C4-2B cells. In VCaP cells, we observed that suppressing either androgen signalling or PAT4 expression, which are functionally linked in these cells, induces a similar exosome switch, providing further support that this mechanism is activated by a range of different stresses and treatments that impact on mTORC1 signalling.

We have previously postulated that because Rab11a-exosomes carry different cargos loaded in recycling endosomal compartments and in HCT116 colorectal cancer cells, they appear to have enhanced pro-tumorigenic effects (Fan et al., 2020), these vesicles may be able to promote adaptation and competition between tumour cells that respond differentially to stresses. In the case of ADT, Rab11a-exosome release might promote growth in those cells that are most resistant to the growth-inhibitory effects of androgen deprivation, for example, if they express high levels of intracellular PAT4. This suggests a model in which these cells might outcompete their neighbours, and therefore lead to increased intracellular PAT4 expression, consistent with our observations in patients. It will now be important inn future studies to assess the activities and bioactive cargos of Rab11a-exosomes from PCa cells, so that this hypothesis can be tested further.

## Materials and Methods

### Electron microscopy

To image isolated EVs, a 5-10 μl drop of EV suspension (~0.2 mg/ml) was adsorbed onto a glow discharged carbon-coated copper grid (300 mesh, 3 mm grid, TAAB Laboratories) for approximately 2 minutes. After adsorption, grids were blotted dry with filter paper and washed with Milli–Q water. To negatively stain the EVs, uranyl acetate (2% w/v) was applied to the grids for 10 seconds, before the excess solution was removed with clean filter paper. The grids were then air dried and imaged on a FEI Tecnai12 TEM (FEI UK Ltd) operating at an accelerating voltage of 120 kV. Images were captured using a bottom-mounted Gatan OneView camera at 120,000x magnification.

### Cell culture

VCaP and C4-2B cells were purchased from ATCC. C4-2B cells were maintained in RPMI-1640 medium (Gibco^®^, Invitrogen) and VCaP cells in high glucose DMEM (Gibco^®^, Invitrogen) supplemented with 10% heat-inactivated fetal bovine serum (FBS, Life Technologies) and 100 U/ml Penicillin-Streptomycin (Life Technologies). All cells were incubated at 37°C with 5% CO_2_ prior to EV collection. DHT was added to 5 nM concentration and 10 μM bicalutamide was used in anti-androgen studies.

### Extracellular vesicle isolation

For EV isolation, cells were plated in complete growth medium at the following densities: 2.5 x 10^7^ VCaP cells per dish and 1 x 10^7^ C4-2B cells per dish in 150 mm dishes (12 dishes/condition). Cells were left to attach overnight and had reached approximately 80% confluency before beginning EV collection. Generally, serum-free media conditions were used when isolating EVs to eliminate contaminating EVs present in FBS. Cells were cultured in standard growth medium supplemented with 1% ITS (Insulin-Transferrin-Selenium; #41400045 Life Technologies) (Tauro et al., 2012). However, for EV isolation from VCaP cells following anti-androgen treatment, medium was supplemented with 1% pre-spun CSS instead of ITS. Briefly, CSS was centrifuged for ~18 hours (100,000 *x g;* 4°C) and the EV-depleted supernatant was filtered through a 0.22 μm pore PVDF Millex-HV filter (Merck Millipore) and analysed by NTA, TEM, and western blotting before being added to media. For glutamine depletion experiments, VCaP cells and C4-2B cells were grown for 24 hours in medium without L-glutamine supplemented with 1% ITS and 2.00 mM or low glutamine (0.02) (Life Technologies). When used, Torin1 was added at 100 nM. For shRNA knockdown of *PAT4*, VCaP cells were cultured for five days following transfection *PAT4* shRNA lentivirus (TRCN0000043984; Sigma). Cells were incubated in the isolation media for a period of 24 hours and EVs were purified from the conditioned media via size-exclusion chromatography (SEC).

After 24 hours, conditioned medium was collected and pre-cleared by centrifugation (500 *x g*, then 2,000 *x g*; 10 minutes each, 4°C) to remove whole cells and cell debris. The supernatant was subsequently vacuum filtered through a 0.22 μm pore membrane (Milex^®^) to eliminate any larger vesicles of non-endosomal origin. The resultant filtrate was concentrated to a volume of approximately 30 ml using the Vivaflow 50 R tangential flow filtration (TFF) system (Sartorius, MWCO = 100 kDa) and further filtered to a final volume of 1 ml by repeated centrifugation (4000 × *g*, 4°C) using an Amicon Ultra-15 spin-filter unit (Merck Millipore, MWCO = 100 kDa). The sample was then loaded onto a 24 ml SEC column packed with sepharose CL-4B, 84 nm pore size (GE Healthcare Life Sciences) set up in an AKTA start system (GE Healthcare Life Science) and eluted as 1 ml fractions in PBS (Gibco^®^, Invitrogen). For EV analysis, Fractions corresponding to the initial “EV peak” (two to seven) were pooled together in 100 kDa Amicon tubes and concentrated to approximately 100 μl via ultrafiltration, then stored at −80°C.

As described by Fan et al (2020), EV preparations for western analysis were assessed after normalisation to the protein mass of EV-secreting cells. For normalisation, following EV isolation, cells were twice washed in PBS and lysed on ice with RIPA buffer supplemented with protease and phosphatase inhibitors (Sigma). Cell debris was pelleted by centrifugation at 13,000g for 10 minutes at 4°C and the protein concentration measured by bicinchoninic acid (BCA) assay (Pierce, Thermo Scientific).

### Nanoparticle tracker analysis (NanoSight^®^)

EV particle size and concentration data were obtained using the NanoSight NS500 instrument (Malvern, UK) equipped with a 405 nm laser, CCD camera and NTA software version 2.3 (NanoSight Ltd.). Samples were diluted in PBS and measured within the linear range of the NS500 between 2-10 × 10^8^ particles/ml. Three videos of 30 seconds were captured between three and five times per EV sample at a known dilution.

### *PAT4* knockdown in VCaP cells

The following conditions were used for *PAT4* knockdown. A total of 5 x 10^5^ HEK293T cells per well were seeded into 6 well plates and allowed to settle overnight. The next day, lentiviral vectors expressing the shRNA targeting *PAT4* were produced as follows: HEK293T cells were co-transfected with shRNA-PAT4 plasmid DNA (5’-CCGGCCTTGATAAATGAGCAGAATTCTCGAGAATTCTGCTCATTTATCAAGGTTT TTG-3’; TRCN0000043984; Sigma) or new shRNA lentivirus (GTTGTCCTTATTGGAGATTC; TRCN0000381243; Sigma), along with psPAX2 packaging and pMD2.G envelope plasmid DNA at a ratio of 4:3:1, respectively. The mixture was incubated at room temperature for 30 minutes and then added to each well in a dropwise manner. Approximately 24 hours post-transfection, the medium was replaced with complete growth media and left to incubate for a further 24 hours prior to the initial collection of the conditioned medium. An additional collection was made after a further 24hours. The conditioned media from the two collections were then combined and cleared by centrifugation (500 *x g*, 4°C, 5 minutes) and filtration through a 0.45 μm pore PVDF Millex-HV filter (Merck Millipore). Finally, the lentiviral media was added to VCaP cell culture along with 8 μg/ml polybrene. The lentiviral medium was incubated for two days, then removed and fresh complete growth medium was added. *PAT4* knockdown was then permitted for three more days prior to changing to fresh EV collection medium and EV isolation.

### Immunohistochemistry

Tissue microarrays (TMAs) were constructed from formalin-fixed paraffin-embedded (FFPE) radical prostatectomy specimens from 137 PCa patients (Gleason grades three and four with negative surgical margins), and from trans-urethral resection of the prostate (TURP) samples from 44 patients with benign prostatic hyperplasia (BPH). The TMA consisted of a single tissue core for each of the 181 patients (accessible cores after accounting for 33 and 20 drop outs respectively; Gonnissen et al., 2017).

TMA sections were obtained from the Oxford Centre for Histopathology Research, Oxford University Hospitals NHS Foundation Trust, Oxford, UK. The TMA contained patient samples from Oxford and was built under the approval of the Oxford Radcliffe Biobank Ethics Committee (reference number 19/SC/0173). The study itself was approved by the ethics committee of KU Leuven, Leuven, Belgium (reference number S55726).

Immunohistochemistry was performed on formalin-fixed, paraffin-embedded TMA PCa and BPH samples. A fully validated anti-PAT4 mouse monoclonal antibody was used to stain the tissue (Fan et al., 2016). PAT4 expression scoring was conducted by an expert uro-pathologist (CV) using the standardised immunoreactive score (IRS) system and presented as the mean ± SEM (Remmele and Stegner, 1987). Separate scores were produced for the presence of PAT4 in the cell cytoplasm and at the cell membrane.

### Quantitative RT–PCR

Total RNA was extracted using the RNAeasy Mini Kit (Qiagen) according to the manufacturer’s protocol. RNA was quantified using the NanoDrop ND −1000 Spectrophotometer (New Brunswick) and then reverse transcribed into cDNA using the high-capacity cDNA Reverse Transcription Kit (Applied Biosystems). Gene expression was assessed via quantitative real-time PCR (qRT-PCR) using Power SYBR Green Master Mix (Applied Biosystems) and monitored using a StepOne^®^ Real-Time PCR system (Applied Biosystems). Each sample was analysed in triplicate and mRNA levels normalised to β-actin.

### Western blot analysis

To prepare samples for western blotting, cells were lysed in RIPA buffer (Sigma) supplemented with a mixture of protease and phosphatase inhibitors, then centrifuged at 13,500 × *g* (10 minutes, 4°C) to remove nuclei and cellular debris. Protein content was quantified using the BCA Protein Assay Kit. To determine the level of mTORC1 signalling phospho-S6 (pS6), assessed relative to total S6 levels, and the γ-phosphorylated form of 4E-BP1, the slowest migrating band detected by a pan-4E-BP1 antibody. Equal amounts of protein, or normalised EV preparations, were dissolved in either reduced (containing 5% β-mercaptoethanol) or nonreduced (for CD63 and CD81 detection) sample buffer and heated (10 minutes, 95°C), then separated on 10% mini PROTEAN^®^ precast gels (BioRad) for 1 hour at 140 V. Separated proteins were transferred to an Immobilin^®^-FL polyvinylidene difluoride (PVDF) membrane (0.22 μm pore size) for 1 hour at 100 V and 4°C. Subsequently, membranes were blocked in 5% skimmed milk in TBS containing 0.1% Tween 20 (TBST) for at least 1 hour, then incubated with primary antibodies diluted in 5% milk in TBST overnight at 4°C. Following incubation, membranes were washed three times in TBST for 5 minutes each, then probed with the relevant HRP-conjugated secondary antibody (diluted in 5% milk in TBST) for 1 hour at RT. The membranes were washed a further three times in TBST for 15 minutes each, and then incubated with Clarity^®^ enhanced chemiluminescence (ECL) solution (Bio-Rad) for 5 minutes before being imaged on the ChemiDoc Touch Imaging System. Relative band intensities were quantified using the ImageJ software. Signals were normalised to tubulin (cell lysates), or CD81 (EVs).

Antibody suppliers, catalogue numbers and concentrations used were: rabbit anti-4E-BP1 (Cell Signaling Technology #9644, 1:1000), rabbit anti-S6 (Cell Signaling Technology #2217, 1:4000), rabbit anti-p-S6-Ser240/244 S6 (Cell Signalling Technology #5364, 1:4000), mouse anti-Tubulin (Sigma #T8328, 1:4000), mouse anti-CD81 (Santa Cruz #23962, 1:500), mouse anti-CD63 (BD Biosciences #556019, 1:500), rabbit anti-Syntenin-1 antibody (Abcam ab133267, 1:500), rabbit anti-Tsg101 (Abcam ab125011, 1:500), mouse anti-Rab11 (BD Biosciences #610657, 1:500), rabbit anti-Calnexin (Abcam #ab213243, 1:1000), mouse anti-GM130 (Novus Biologicals H00002801-B01P), anti-mouse IgG (H+L) HRP conjugate (Promega #W4021, 1:10000), anti-rabbit IgG (H+L) HRP conjugate (Promega #W4011, 1:10000), mouse anti-Alix (Abcam #ab117600, 1:500), rabbit anti-LAMP-1 (Cell Signalling #9091S, 1:1000), rabbit anti-EEA1 (Cell Signalling #3288S, 1:500), rabbit anti-Flotillin-2 (Cell Signalling #3436S, 1:500), mouse anti-PAT4 (Novus #NBP2-50237, 1:200)

### Growth assay

For growth assays, VCaP cells were transduced with shNT or shPAT4 for 5 days then plated in basal culture medium supplemented with 10% charcoal-stripped serum (CSS; Gibco^®^, Invitrogen) and left to attach overnight. The following day, cells were treated with DHT (0.1 nM) or with equal volumes of vehicle (MeOH) for 48 hours. Phase-contrast images were captured every two hours using the IncuCyte ZOOM^®^ (10X magnification; Essen Bioscience) and cell growth was analysed using the IncuCyte^®^ software to automatically detect cell edges and to generate a confluence mask for cell coverage area calculation. Each biological replicate is represented as mean of percentage confluency for at least eight technical replicates. All assays were repeated on at least three independent occasions.

### Statistical analysis

Unless otherwise stated, data are presented as the mean ± SD. A two-tailed Student’s t-test was used to compare differences between subgroups. A statistically significant difference was defined as \**P*<0.05, \*\**P* < 0.01 or \*\*\**P* < 0.001.

## Acknowledgements

We are particularly grateful to Daniel Stevens for his contributions to the analysis of the patient samples and the early aspects of this work, and to Clive Wilson for advice during this study. We thank Errin Johnson and the Dunn School EM Facility for help with the EM analysis. We acknowledge the Oxford Radcliffe Biobank and OCHRe for provision and approvals required to build the patient TMAs.

## Supplementary Figures

**Supplementary Figure 1.**
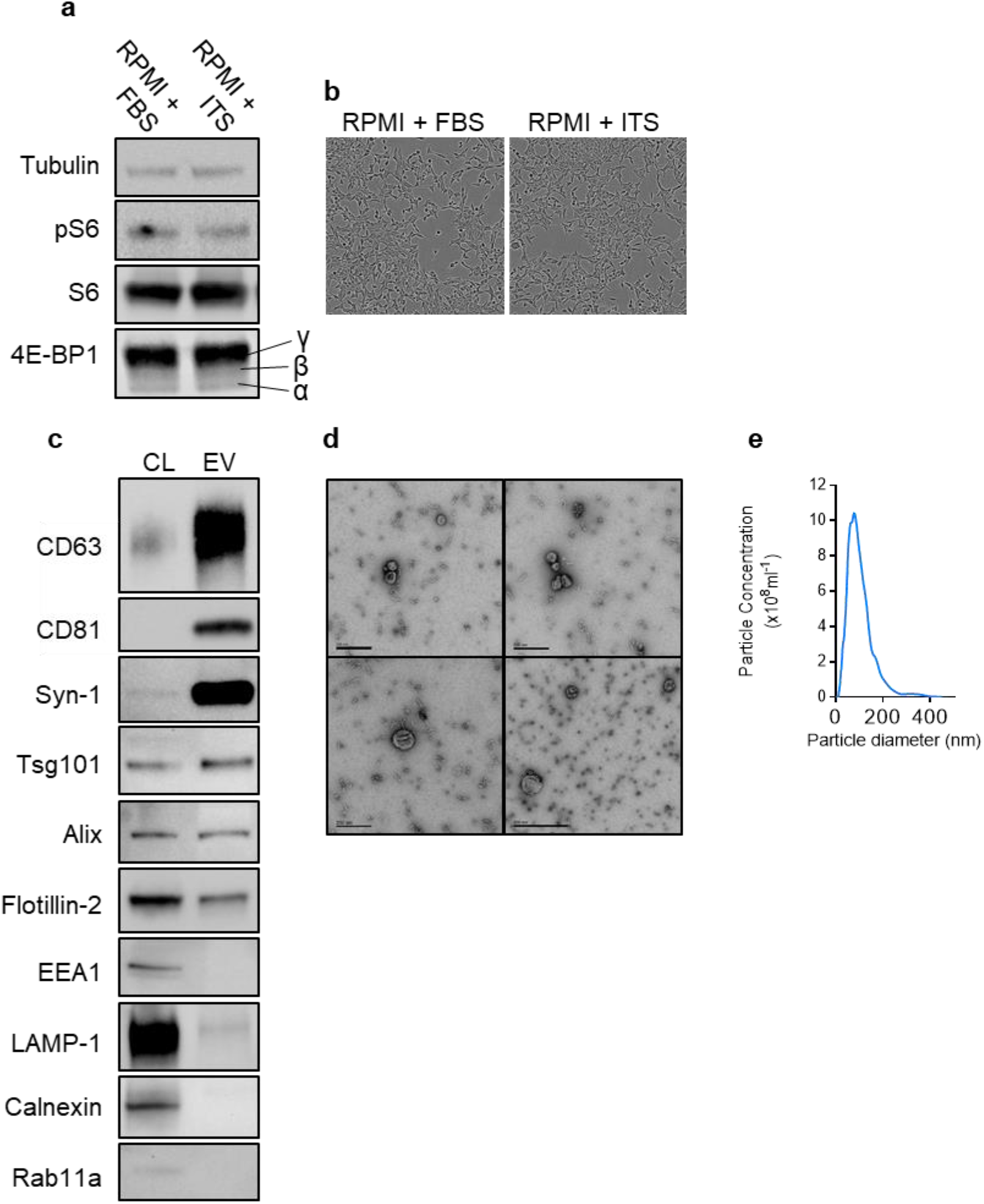
Characterisation of EV isolation conditions for C4-2B cells. (**a**,) Culture conditions for collecting EVs from C4-2B cells were determined by testing different media for maintenance of mTORC1 activity during the collection period. (**b**) Cell confluency was not altered by culture in the two media. (**c,d**) EVs were characterised using (**c**) western blot analysis, (**d**) transmission electron microscopy (TEM) and (**e**) NanoSight Tracking Analysis (NTA).

**Supplementary Figure 2.**
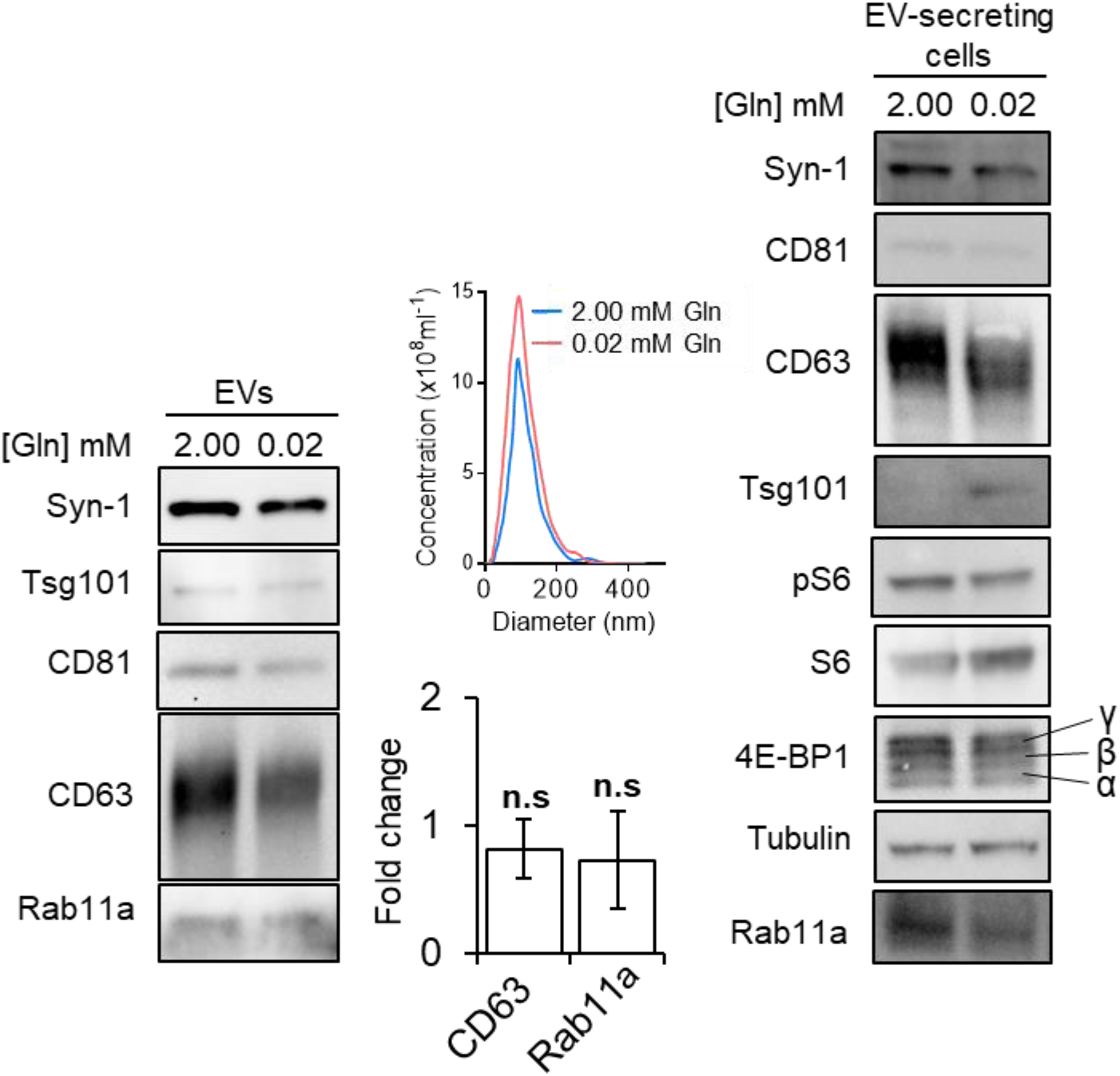
Glutamine starvation of C4-2B cells does not induce Rab11a-exosome secretion. Western blot analysis of cell lysates and EVs from C4-2B cells in glutamine-replete (2.00 mM) and glutamine-depleted (0.02 mM) medium for 24 h. The activity of mTORC1 was assessed via phosphorylation of S6 (pS6) and 4E-BP1. EV lysates were probed for putative exosome proteins. Bar charts present data from three biological repeats. The number of EVs in each condition was determined using NanoSight analysis. ns > 0.05

**Supplementary Figure 3.**
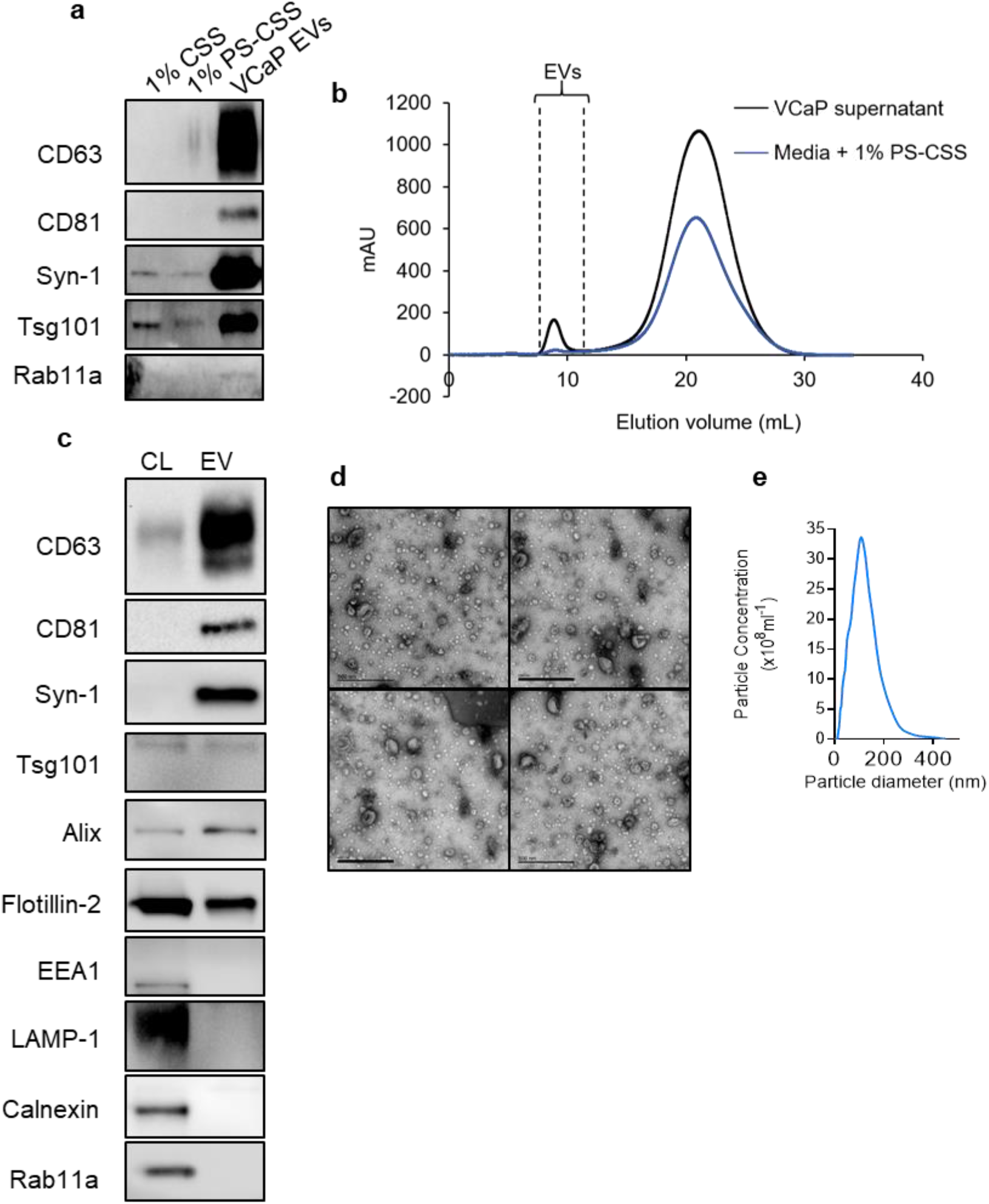
Characterisation of EV isolation conditions for VCaP cells using pre-centrifuged, charcoal-stripped serum. To assess the effect of bicalutamide on VCaP cells, alternative conditions for culturing these cells were developed using 1% pre-spun, charcoal-stripped serum (PS-CSS). (**a**) Western blot analysis revealed a negligible contribution of exosome proteins from serum EVs either before (CSS) or after (PS-CSS) centrifugation. (**b**) EVs isolated by this method were further characterised using (**c**) western analysis, (**d**) transmission electron microscopy (TEM) and (**e**) NanoSight Tracking Analysis (NTA).

